# Dysregulation of placental mitochondrial structure dynamics and clearance in maternal obesity and gestational diabetes

**DOI:** 10.1101/2025.07.17.665416

**Authors:** Leena Kadam, Kaylee Chan, Ethan Tawater, Leslie Myatt

## Abstract

**Introduction:** The placenta is exposed to an altered metabolic environment in obesity and gestational diabetes (GDM) leading to disruption in placental function. Mitochondria are critical for energy production and cellular adaptation to stress. We previously reported reduced trophoblast mitochondrial respiration in GDM. Here we examine changes in mitochondrial structure dynamics, quality and protein homeostasis as well as clearance in both obese and GDM placentas of male and female fetuses. As obesity significantly increases the risk for GDM, our goal is to determine the distinct effects of each on placental mitochondria.

**Methods:** We collected placental villous tissue following elective cesarean section at term from lean (LN, pre-pregnancy BMI 18.5-24.9), obese (OB, BMI>30) or obese with type A2 GDM women. Expression of proteins involved in mitochondrial biogenesis, structure dynamics, quality control and clearance were assessed by Western blotting. Significant changes between groups were determined in fetal sex-dependent and independent manner.

**Results:** Only placentas from obese women showed increase in proteins regulating mitochondrial biogenesis (PGC-1α and SIRT1). We report fetal sex-specific changes in mitochondrial fusion but an overall decline in fission in OB and GDM placentas. Both maternal obesity and GDM affected proteins involved in maintaining mitochondrial protein quality and genome stability. This was accompanied by a reduction in mitochondrial complexes, suggesting impaired mitochondrial function. Obesity led to partial activation of mitophagy pathways (e.g., increased PINK1 without PARKIN activation), GDM placentas failed to mount this response.

**Discussion:** Obesity and GDM affect placental mitochondria through distinct complex sex-specific mechanisms that may contribute to altered mitochondrial function.

## INTRODUCTION

The incidence of obesity (BMI>30) is increasing significantly worldwide with the WHO describing this obesity “epidemic” as an escalating threat to population health and healthcare systems. One third of women in the United States are obese and one half of pregnant women are either overweight (BMI 25.0-29.9) or obese [1]. Obese women face a higher risk for spontaneous abortion, recurrent miscarriage, and stillbirth [2-4]. Pre-pregnancy obesity also increases the risk of developing gestational diabetes mellitus (GDM)[5-8] by 3 fold[9, 10]. GDM is classified as “diabetes first diagnosed in the 2^nd^/3^rd^ trimester of pregnancy that is clearly neither preexisting type 1 or type 2 diabetes (T2DM)” and is estimated to occur in ∼7-10% of pregnancies worldwide[11-13]. Women with obesity and GDM display hyperglycemia and hyperlipidemia[14], are more susceptible to hypertensive disorders, dysfunctional labor, cesarean delivery, thromboembolic events [1, 15, 16] and are at increased risk of developing type 2 diabetes in later life [17]. Fetal complications in these pregnancies include congenital malformations, large-for-gestational-age (LGA) infants, intrauterine growth restriction (IUGR), asphyxia and stillbirth. Importantly fetal programming for subsequent development of obesity and diabetes [18-20] [1, 9, 15, 21, 22] is a feature of obese and GDM pregnancies contributing to the vicious cycle of their development. Women carrying a male fetus have been reported to be at a higher risk for developing GDM and of a more severe nature and suffering more adverse pregnancy outcomes underscoring the sexual dimorphic effects of these conditions [23, 24].

Our work focuses on understanding placental adaptations in obese and GDM pregnancies. The primary function of the placenta is to regulate maternal metabolism to ensure optimal fetal growth and development. In pregnancies complicated by obesity and GDM, the placenta is exposed to a hyperglycemic and hyperlipidemia environment. Our previous studies have shown that placental trophoblast isolated from such pregnancies exhibits mitochondrial dysfunction including changes in respiration (oxidative phosphorylation)[25], altered metabolic flexibility in utilizing different substrates (glucose vs glutamine vs fatty acids) for ATP production via the TCA cycle as well as reduction in expression of mitochondrial complexes I-V and increased oxidative stress[26, 27]. These changes were sexually dimorphic in nature. Mitochondria are central to cellular energy metabolism and their dysfunction is implicated in several diseases including obesity mediated adipose tissue dysfunction and diabetes[28-32]. They are also highly dynamic organelles which undergo biogenesis, structural changes and degradation in response to cellular stress to ensure cell survival and function. Here we aim to understand changes in placental mitochondrial dynamics in response to the maternal metabolic stress of obesity and GDM. We also evaluate if these changes are influenced by fetal sex.

## METHODS

### Sample collection

Informed consent was obtained from all patients under a protocol approved by the Institutional Review Board of Oregon Health & Science University, prior to tissue being de-identified and placed in a tissue repository. Maternal blood and placental tissue were collected from three groups of women: Lean (LN, pre-pregnancy BMI 18.5-25.0), Obese (OB, pre-pregnancy BMI 30.1-45.0) and obese with type A2GDM (OB+GDM) (BMI matched to obese group) with either a male or a female fetus (n=12 each group and n=6 per sex). GDM was defined using IADPSG criteria which recommends 75-grams two hours OGTT with at least one abnormal result: Fasting plasma glucose (FPG) at ≥ 92 mg/dl or 1-hour ≥ 180 mg/dl or 2-hour ≥ 153mg/dl to be classified as GDM[33]. Women with type A2 GDM were those with greater than 30% of FPG ≥90-95 mg/dl or 1 hr Postprandial ≥130-140 mg/dl and needed medication to control blood glucose. For consistency, we only collected samples from women who received insulin as medication. We acknowledge that women with pre-pregnancy BMI in the lean range also develop GDM, however obese women have 3 times higher risk for developing GDM hence we focused on this at-risk group [9, 10]. Inclusion criteria: a singleton pregnancy, an age range of 18-45 and delivery by elective cesarean section at term with no labor. Exclusion criteria: concurrent disease (including, but not limited to IUGR, hypertension, pre-eclampsia, eating disorders, infection, inflammatory disorders), use of tobacco, drugs, or medications other than to treat GDM, excessive weight gain or loss prior to pregnancy (>20 lbs), or bariatric surgery in the last year and labor defined by regular uterine contractions (every 3-4 minutes, verified by tocodynamometry) resulting in cervical dilatation and/or effacement. Placental villous tissue was immediately sampled at delivery from 5 random sites, flash frozen and stored at -80°C. Subsequently villous tissue was powdered under liquid nitrogen and equal amounts of the 5 separate samples combined before analysis for each subject.

### Protein extraction and Western blotting

Approximately 25mg of powdered tissue was mixed with 250μL of RIPA buffer supplemented with protease/phosphatase inhibitors (ThermoFisher Scientific, Cat. #A32959) and incubated on ice for 30 minutes. Post incubation, the tubes were centrifuged at 1000 rpm for 5 minutes and supernatant/lysate transferred to a new tube. Total protein was quantified using the Pierce BCA Protein Assay Kit per manufacturers protocol. (ThermoFisher Scientific, Cat. #23225). Approximately 30μg of protein was separated on 12% sodium dodecyl sulphate-polyacrylamide gel electrophoresis (SDS-PAGE) hand-cast gels for approximately 2 hrs at 100V and transferred onto nitrocellulose membranes using Mini-PROTEAN tetra cell electrophoresis chamber (BioRad, Cat. # 1658004). Membranes were blocked in 5% (w/v) nonfat milk in TBS + 0.1% Tween 20 (TBST) for 1 hr and incubated with primary antibodies (see supplementary table 1) overnight at 4 °C. Next day the membranes were washed three times in TBST for 5 min each and incubated with HRP-conjugated secondary antibodies. Membranes were washed and the antibody binding was detected using Supersignal West Pico Plus ECL Substrate (ThermoFisher Scientific, Cat. #34578) for 5 min and imaged using ChemiDoc Imaging System (Bio-Rad Laboratories, USA) and Image Lab V.5.1 software (Bio-Rad Laboratories, USA). Densities of immunoreactive bands were measured as arbitrary units by ImageJ software. Protein levels were normalized to a housekeeping protein β-actin (1: 20,000; Abcam, UK). To assess phosphorylated and total (unphosphorylated and phosphorylated) proteins, the membranes were first probed with phospho-antibodies, then stripped using mild stripping buffer (Thermo Scientific™ Restore™ Fluorescent Western Blot Stripping Buffer). The stripped blots were then blocked again as outlined above and probed with the antibody detecting total protein.

### Statistical analysis

Data was analyzed for statistical difference using the GraphPad Prism 7.0 software. All analyses consisted of n=6 samples per sex per clinical group. Outliers were detected using the ROUT method and wherever detected maximum 2 outliers were excluded from analysis. The values between groups were analyzed using ANOVA followed by post hoc Tukey analysis.

## RESULTS

### I. Clinical characteristics

The clinical characteristics of patients for whom placental tissue was utilized are outlined in Table I. There was no difference in maternal age and gestational age between LN, OB and OB+GDM groups. As expected, patients from the OB and OB+GDM group had significantly higher BMI compared to LN. All neonates had good outcomes, were born appropriate for gestational age and there were no significant differences in fetal and placental weights and fetal: placental weight ratio between the clinical groups.

**Table 1:** Clinical characteristics of study participants. Data presented as Mean ± SD. Significant differences were determined using ANOVA. * *p* < 0.05 for comparisons with the LN group. ^ϯ^ p<0.05 for same sex comparisons with the LN group and †*p* < 0.05 for male vs female comparisons within the same clinical group.

|  | Lean<br>(LN, n=10) |  | Obese<br>(OB, n=10) |  | Gestational Diabetes<br>(GDM, n=10) |  |
| --- | --- | --- | --- | --- | --- | --- |
|  | Male<br>(n=6) | Male<br>(n=6) | Male<br>(n=6) | Male<br>(n=6) | Male<br>(n=6) | Female<br>(n=6) |
| Maternal age (yrs) | 35.9 ± 4.4 |  | 32.4 ± 5.5 |  | 33.0 ± 3.5 |  |
|  | 35.5 ± 3.5 | 36.3 ± 5.5 | 33.0 ± 5.9 | 31.8 ± 5.6 | 32.3 ± 4.1 | 33.8 ± 2.9 |
| Maternal BMI<br>(wt[kg]/height[m<sup>2</sup>]) | 20.9 ± 1.3 |  | 37.9 ± 8.0* |  | 40.2 ± 10.8* |  |
|  | 20.6 ± 1.2 | 21.1 ± 1.4 | 37.7 ± 10.1<sup>†</sup> | 38.0 ± 6.1<sup>†</sup> | 41.7 ± 12.7<sup>†</sup> | 38.7 ± 9.4<sup>†</sup> |
| Gestational age<br>(wks) | 38.6 ± 0.8 |  | 37.63 ± 1.1 |  | 38.07 ± 1.0 |  |
|  | 39.0 ± 0.08 | 38.1 ± 1.0 | 38.4 ± 0.9 | 36.8 ± 0.5<sup>†</sup> | 37.9 ± 0.9 | 38.1 ± 1.2 |
| Fetal birthweight<br>(g) | 3452 ± 364.5 |  | 3327 ± 335.3 |  | 3297 ± 715.5 |  |
|  | 3672 ± 301 | 3233 ± 292 | 3425 ± 340 | 3228 ± 328 | 3687 ± 534 | 2906 ± 688 |
| Placental weight<br>(g) | 491.1 ± 91.84 |  | 604.7 ± 129.9 |  | 572.2 ± 131.2 |  |
|  | 479.8 ± 116.4 | 504.1 ± 68.2 | 574.6 ± 140.6 | 640.8 ± 120.5 | 635.4 ± 155.2 | 519.5 ± 88.4 |
| Fetal:Placental<br>weight ratio | 6.7 ± 0.6 |  | 5.71 ± 1.23 |  | 5.74 ± 1.21 |  |
|  | 7.0 ± 0.4 | 6.4 ± 0.6 | 6.2 ± 1.3 | 5.1 ± 0.7 | 5.8 ± 1.3 | 5.6 ± 1.2 |

### II. Maternal obesity alters mitochondrial biogenesis in placenta

We studied expression of Sirtuin 1 (SIRT1) and Peroxisome proliferator-activated receptor-γ coactivator-1 alpha (PGC1α) as markers for mitochondrial biogenesis. We did not observe any statistically significant changes in SIRT1 levels, however, the OB group showed high variability (Fig 1A). Expression of PGC1α was significantly higher in the OB group compared to both LN and OB+GDM groups (Fig 1C). Fetal sex-stratified analysis for SIRT1 showed similar expression levels in male and female placentas from LN and OB+GDM groups. Both male and female placentas from the OB group also had similar levels with high variability (Fig 1B). Fetal sex-stratified analysis for PGC1α showed comparable expression in male and female placentas from LN and OB+GDM groups. However, in the OB group, females had significantly higher expression vs the males as well as vs females from LN and OB+GDM groups (Fig 1D). Higher levels of PGC1α suggest an increase in signals for mitochondrial biogenesis in the OB group.

**Figure 1:**
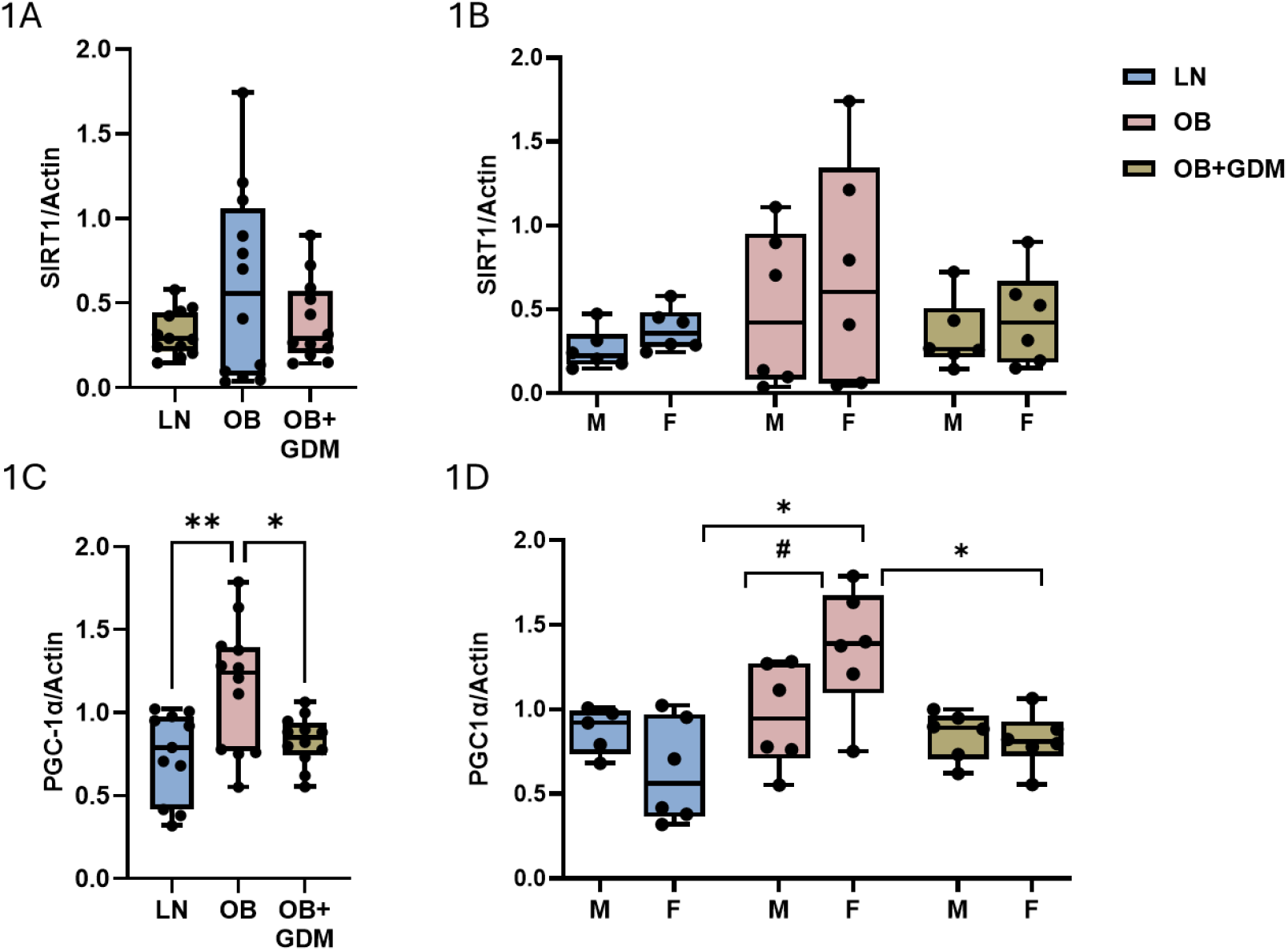
Expression of mitochondrial biogenesis related proteins in villous tissue from LN, OB and OB+GDM placentas. A. B. Expression of SIRT1 and PGC-1α in fetal sex combined and C, D in a sex-stratified manner respectively. Data presented as box and whispers plot showing median, interquartile range, minimum and maximum values. * p<0.05, **p<0.01 and #p<0.05 for comparison between male and female.

### III. Mitochondrial fusion-fission dynamics are altered in placentae from obese and GDM pregnancies

We evaluated the expression of Optic Atrophy 1 (OPA1) a mitochondrial dynamin GTPase-like protein which regulates mitochondrial membrane fusion and Dynamin-related protein 1 (DRP1) which initiates membrane fission. Expression levels of OPA1 were higher although not statistically significant in OB+GDM placentas when compared to LN and OB placentas (Fig 2A). Fetal sex-stratified analysis however revealed a sexually dimorphic pattern with significantly higher levels in female placentas from LN and OB compared to their respective male counterparts. This dimorphism was absent in the OB+GDM group where the male and female placentas had similar expression levels. Furthermore, male placentas from the OB+GDM group had significantly higher expression compared to LN and OB males, but this trend was absent in female placentas (Fig 2B). To initiate fission, DRP1 needs to be activated by phosphorylation at Ser-616 whereas phosphorylation at other serine residues leads to its inactivation[34]. We assessed levels of both phosphorylated Ser-616 DRP1 and total DRP1 and calculated the ratio of phosphorylated: total DRP1 to determine DRP1 activation. We observed a significant decline in levels of activated DRP1 in OB and OB+GDM placentas compared to the LN group (Fig 2C). Fetal sex stratified analysis revealed that this decrease could be accounted for by a decrease in male placentas only with no significant change in female placentas (Fig 2D). The levels of total DRP1 were comparable across the 3 groups in the sex combined analysis, but in the sex-stratified analysis female placentas showed lower levels (not significant) of DRP1 compared to their respective males which showed much greater variability in levels. (Supplemental Fig 1A). Levels of Ser-616 phosphorylated DRP1 (normalized to Actin) also did not show any differences in the sex combined analysis. However similar to total DRP, female placentas showed lower levels of Ser-616 DRP1 compared to the more variable levels in their respective males in all 3 clinical groups (Supplemental Fig 1B). Taken together, these results suggested that as maternal metabolic state worsens from obesity to GDM, mitochondrial fission is significantly reduced and fusion significantly increased in male placentas whereas fission is low in all three metabolic states in female placenta with a corresponding increase in fusion, highlighting sexual dimorphism in the fission/fusion response that impacts the effect of obesity and GDM.

**Figure 2:**
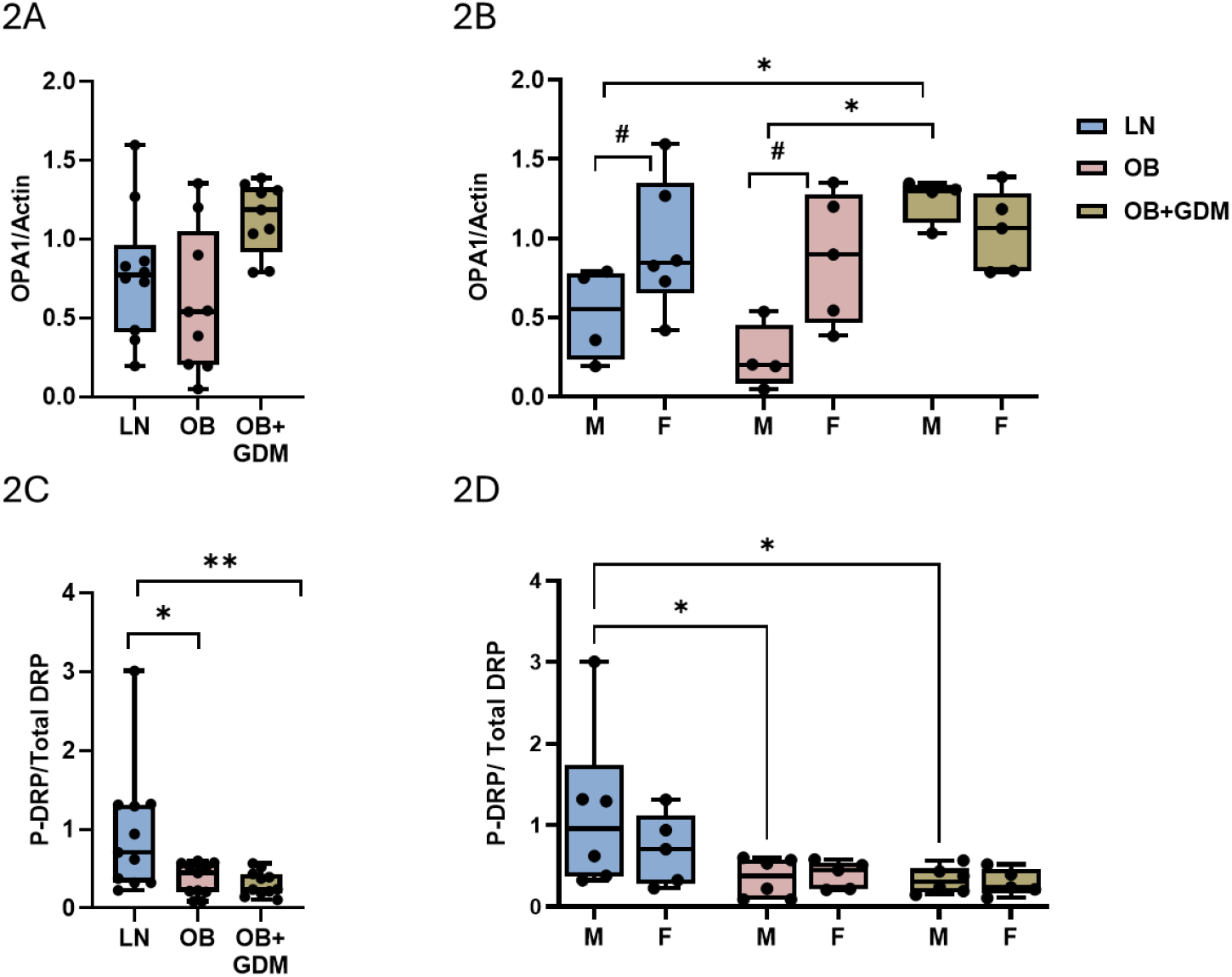
Expression of mitochondrial fusion and fission related proteins in villous tissue from LN, OB and OB+GDM placentas. A. B. Expression of OPA1 and activated DRP1 in fetal sex combined and C, D in sex-stratified manner respectively. Data presented as box and whispers plot showing median, interquartile range, minimum and maximum values. * p<0.05, **p<0.01 and ^#^p<0.05 for comparison between male and female

### IV. Placental mitochondrial genome integrity and protein turnover is altered by maternal obesity and GDM

We assessed the levels of two proteins involved in maintaining mitochondrial quality and genome integrity: (i) Transcription Factor A, Mitochondrial (TFAM) which is essential for transcription, replication, and packaging of mtDNA into nucleoids, thus maintaining mtDNA genome integrity[35] (ii) Lon Protease 1 (LONP1) - a mitochondrial matrix protease that maintains mitochondrial protein homeostasis (by degrading misfolded/damaged proteins) and also participates in regulation of mitochondrial gene expression [36].

We did not see any changes in expression of TFAM across the clinical groups (Fig 3A). Fetal sex-stratified analysis showed that females from all three groups had higher expression compared to the respective males, however these differences were not statistically significant (Fig 3B). In contrast, we observed a significant decline in placental LONP1 expression as maternal condition worsened from obesity to GDM (Fig 3C). Fetal sex-stratified analysis showed no differences between placentas from male and female fetuses in any clinical group (Fig 3D). However, male placentas from OB and OB+GDM groups had significantly lower levels compared to those from the LN group. A similar pattern was observed while comparing the female placentas from GDM group, but the differences were statistically significant only for the OB+GDM vs LN comparison. The absence of change in TFAM expression suggests no impact on mtDNA replication. However, the decrease in LONP1 expression with obesity and GDM suggests dysregulation of mitochondrial gene expression and protein homeostasis[35, 36].

**Figure 3:**
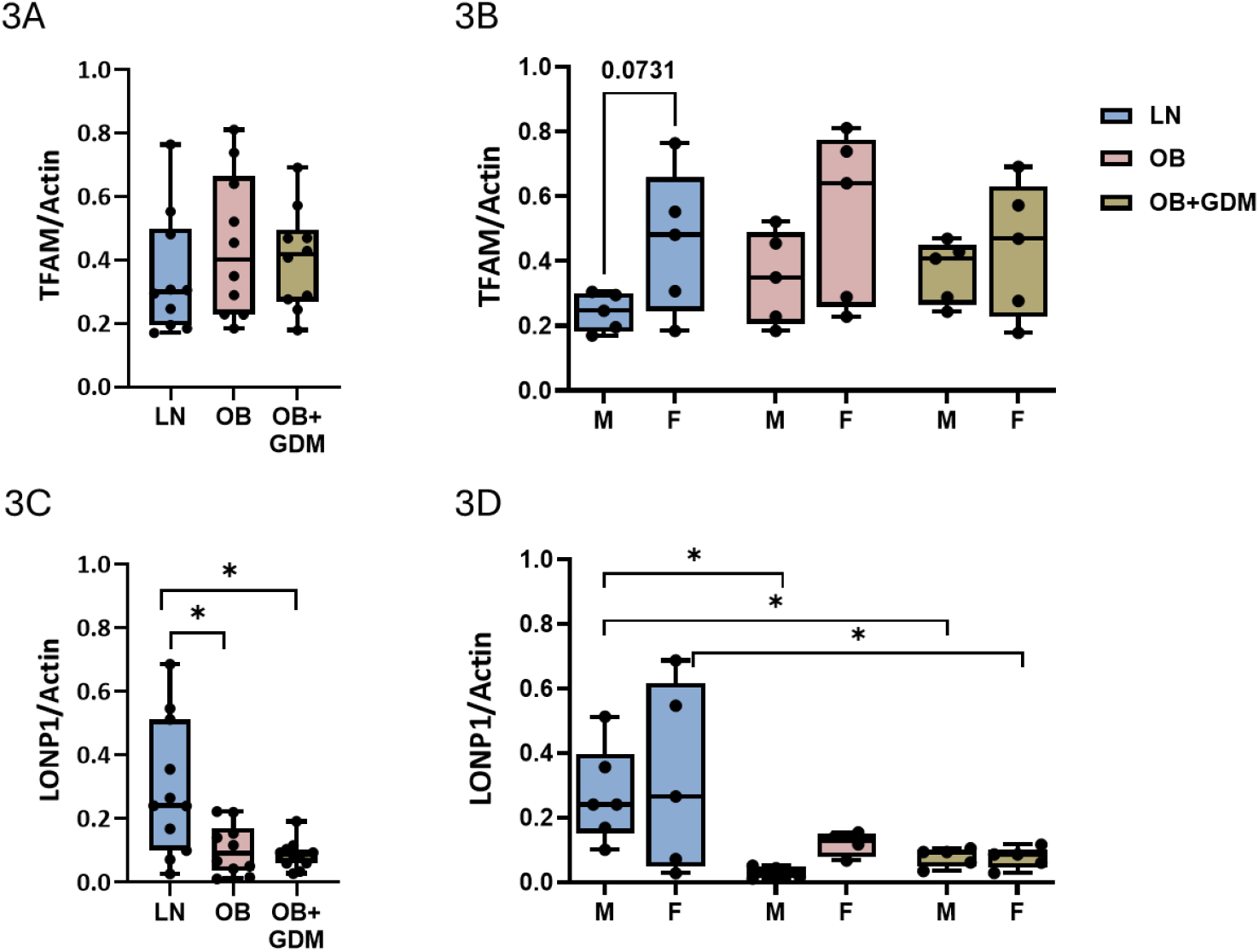
Expression of mitochondrial quality maintenance related proteins in villous tissue from LN, OB and OB+GDM placentas. A. B. Expression of TFAM and LONP1 in fetal sex combined and C, D in sex-stratified manner respectively. Data presented as box and whispers plot showing median, interquartile range, minimum and maximum values. * p<0.05.

### V. Maternal obesity and GDM affect placental mitochondrial content

There are several ways to quantify mitochondrial content with mtDNA, recently shown to correlate to TFAM protein levels,[37, 38] being the most commonly used. As we did not see changes in TFAM levels we did not expect to see changes in mtDNA. Therefore we quantified protein levels of complexes I-V in the Electron Transport Chain (ETC) that create the proton gradient by oxidative phosphorylation to generate ATP [39, 40] to estimate mitochondrial content.

In the sex combined analysis, Complex I (NADH: ubiquinone oxidoreductase) levels were comparable across the three clinical groups (Fig 4A). We observed that complexes II to V had a trend to decrease as maternal condition worsened from obesity to GDM with complex V being significantly downregulated in the OB+GDM group compared to LN (Fig 4B-E). Complex V consists of ATP synthase and reduced levels in OB+GDM could be interpreted as reduced capacity for ATP production in these placentas. We did not observe any fetal sex-based differences in Complex I and V levels (Fig 4F, J). For complex II (succinate dehydrogenase), females from the OB+GDM group had significantly lower levels compared to females from the LN group, but no such differences were observed in the males (Fig 4G). For complex III (ubiquinol–cytochrome coxidoreductase), females from OB and OB+GDM group had significantly lower levels compared to their respective males while no such difference was observed in the LN group (Fig 4H). Complex IV (cytochrome c oxidase) levels were significantly higher in females from LN group compared to their male counterparts as well as females from OB and OB+GDM groups (Fig 4I). Complex IV consists of cytochrome C oxidase which is the rate limiting step of the ETC. Increased levels in LN females (vs LN-male) suggests an inherent difference in ETC activity between male and female placentas. The reduced levels of complex II, III and IV only in females from OB and OB+GDM groups highlight how male and female placentas respond differentially to the maternal metabolic environment. Reduced expression of these enzymes coupled with altered mitochondrial structure could contribute to the mitochondrial dysfunction reported in GDM placentas.

**Figure 4:**
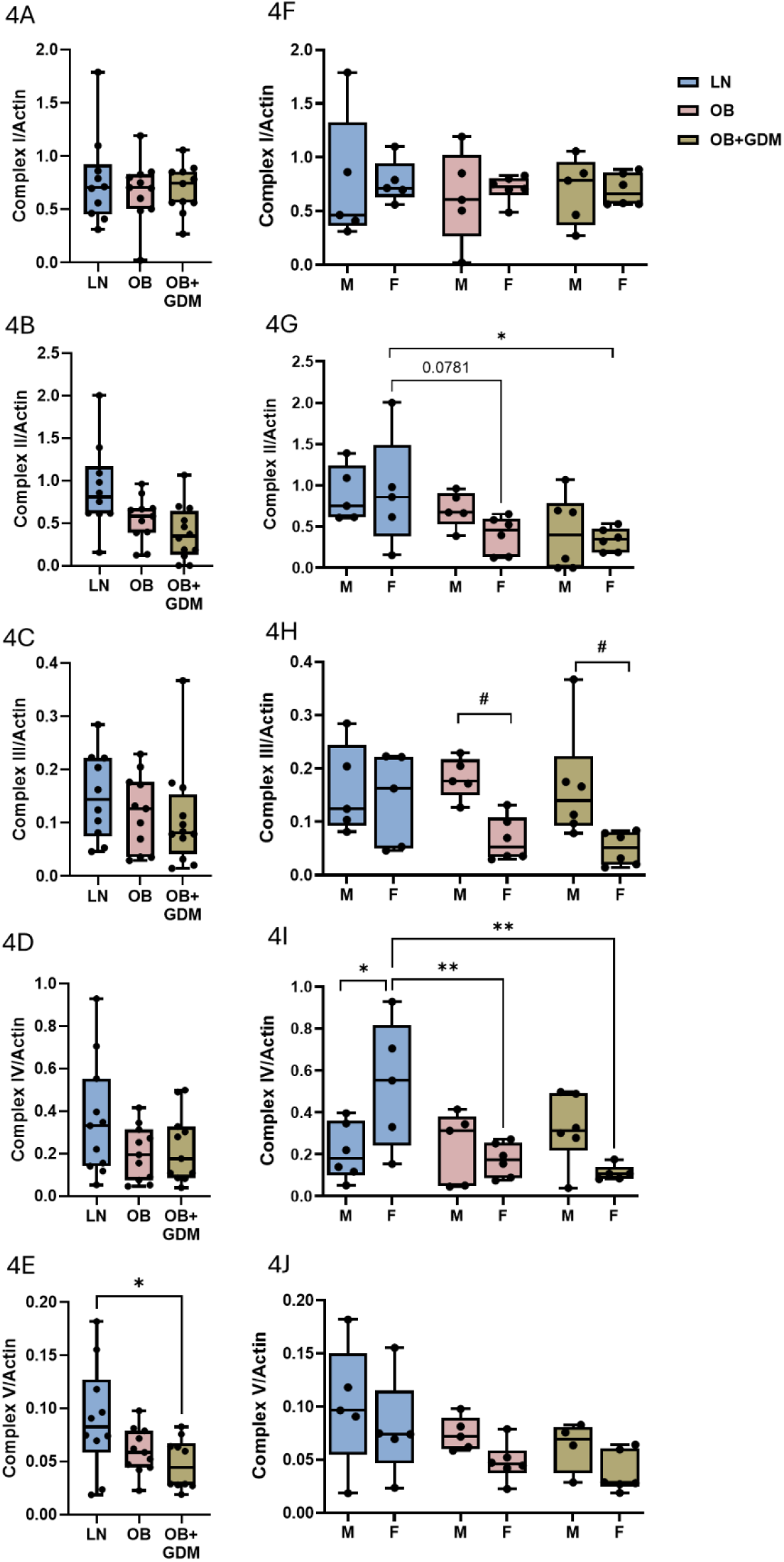
Expression of mitochondrial complexes I-V in villous tissue from LN, OB and OB+GDM placentas. A-E. Expression of complexes I-V in fetal sex combined and F-J in sex-stratified manner respectively. Data presented as box and whispers plot showing median, interquartile range, minimum and maximum values. * p<0.05, **p<0.01 01 and #p<0.05 for comparison between male and female.

### VI. Maternal obesity and GDM alter placental expression of autophagy pathway mediators

As we observed changes in proteins involved in regulating mitochondrial structure dynamics with obesity and GDM, we wanted to assess if there are any effects on mitophagy. Mitophagy is a type of macro autophagy aimed at mitigating stress by eliminating damaged mitochondria from the cells. The protein PINK1 accumulates on the outer membrane of damaged mitochondria and recruits PARKIN to the mitochondrial surface. It also activates PARKIN by phosphorylation at site Ser65 to initiate the mitophagy process. PARKIN is an E3 ubiquitin ligase which causes ubiquitination of other mitochondrial proteins and recruitment of downstream autophagy mediators like LC3 and p62.

We observed that expression of PINK1 significantly increased in placentas from OB group compared to both LN and OB+GDM groups (Fig 5A). Fetal sex-stratified analysis showed that females from the LN and OB groups had significantly lower levels compared to their respective males, but this was absent in the OB+GDM group (Fig 5B). Male placentas from the OB group had significantly higher expression compared to males from both LN and OB+GDM groups. We observed no differences in proportion of activated PARKIN across the clinical groups (Fig 5C). Fetal sex-stratified analysis showed that females from the LN and OB groups had higher expression compared to their respective males, but this sexual dimorphism was absent in the OB+GDM group (Fig 5D). Similar observations were made for levels of phosphorylated PARKIN (normalized to Actin) wherein there were no changes in the sex combined analysis, but fetal sex-stratified data showed higher (statistically non-significant) levels in female placentas vs their male counterparts in all 3 clinical groups (Supplementary Fig 2A). Levels of total PARKIN (normalized to Actin) did not show any changes in fetal sex-combined or stratified analysis (Supplementary Fig 2B). Increases in placental PINK1 expression but no downstream change in PARKIN activation suggests dysregulation of the PINK1-PARKIN axis in obesity. Furthermore, since OB+GDM group showed no changes, it implies that maternal GDM has a distinct influence on placental mitophagy.

**Figure 5:**
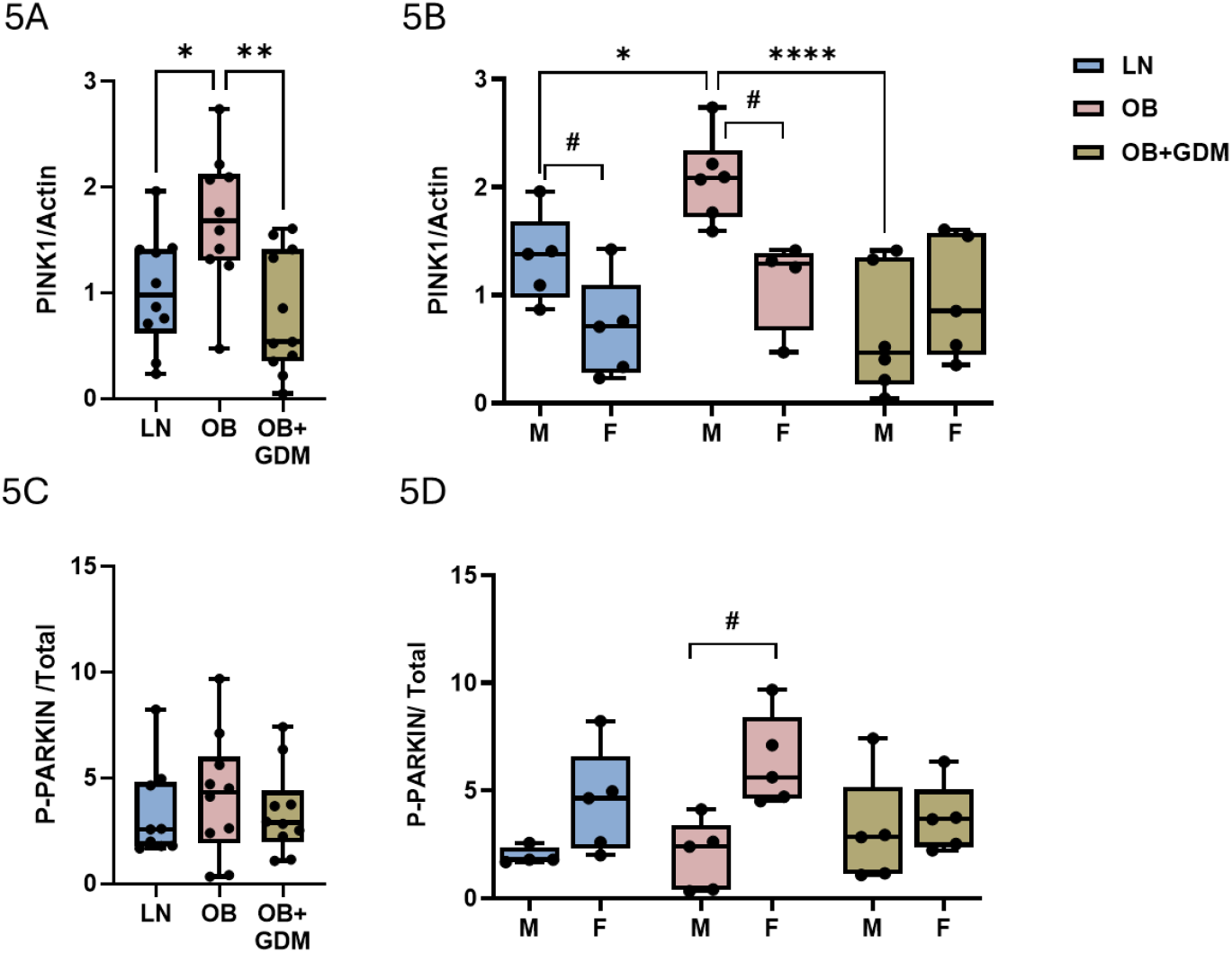
Expression of mitophagy related proteins in villous tissue from LN, OB and OB+GDM placentas. A. B. Expression of PINK1 and activated PARKIN in fetal sex combined and C, D in sex-stratified manner respectively. Data presented as box and whispers plot showing median, interquartile range, minimum and maximum values. * p<0.05, **p<0.01, ****p<0.001 and #p<0.05 for comparison between male and female.

We also analyzed levels of LC3 and p62 autophagy markers to assess if the changes in mitophagy specific proteins are due to dysregulation of overall placental autophagy. LC3 is lipidated to form the active LC3-II form which marks the surfaces of autophagosomes and as such is an indicator of autophagosome formation. A ratio of LC3-II to I was calculated to determine autophagic flux and we observed that OB+GDM placentas had significantly higher ratio compared to both LN and OB placentas suggesting increased autophagic flux in GDM (Fig 6A). Fetal sex-stratified analysis did not show any sexually dimorphic trends, but the ratio was significantly higher in female placentas from OB+GDM group compared to their LN and OB counterparts (Fig 6B). Levels of p62 declined, as maternal metabolic state worsened from obesity to GDM but were statistically significant only for the LN vs OB+GDM comparison (Fig 6C). Fetal sex-stratified analysis showed that while male and female placentas from the LN group had similar levels, female placentas from OB and OB+GDM group had significantly lower levels compared to their respective males (Fig 6D). We also observed high variability in p62 expression in LN group. P62 and LC3 can be used together as a combined marker to assess autophagy levels, with a high LC3-II: LC3-I ratio and low p62 levels indicative of activated and intact autophagy processes[41]. Our results thus indicate an increase in overall autophagy in OB+GDM placentas. Combined with the mitophagy-related PINK1 and PARKIN data, we propose that metabolic stress in GDM increases overall cellular autophagy with no effects on mitophagy.

**Figure 6:**
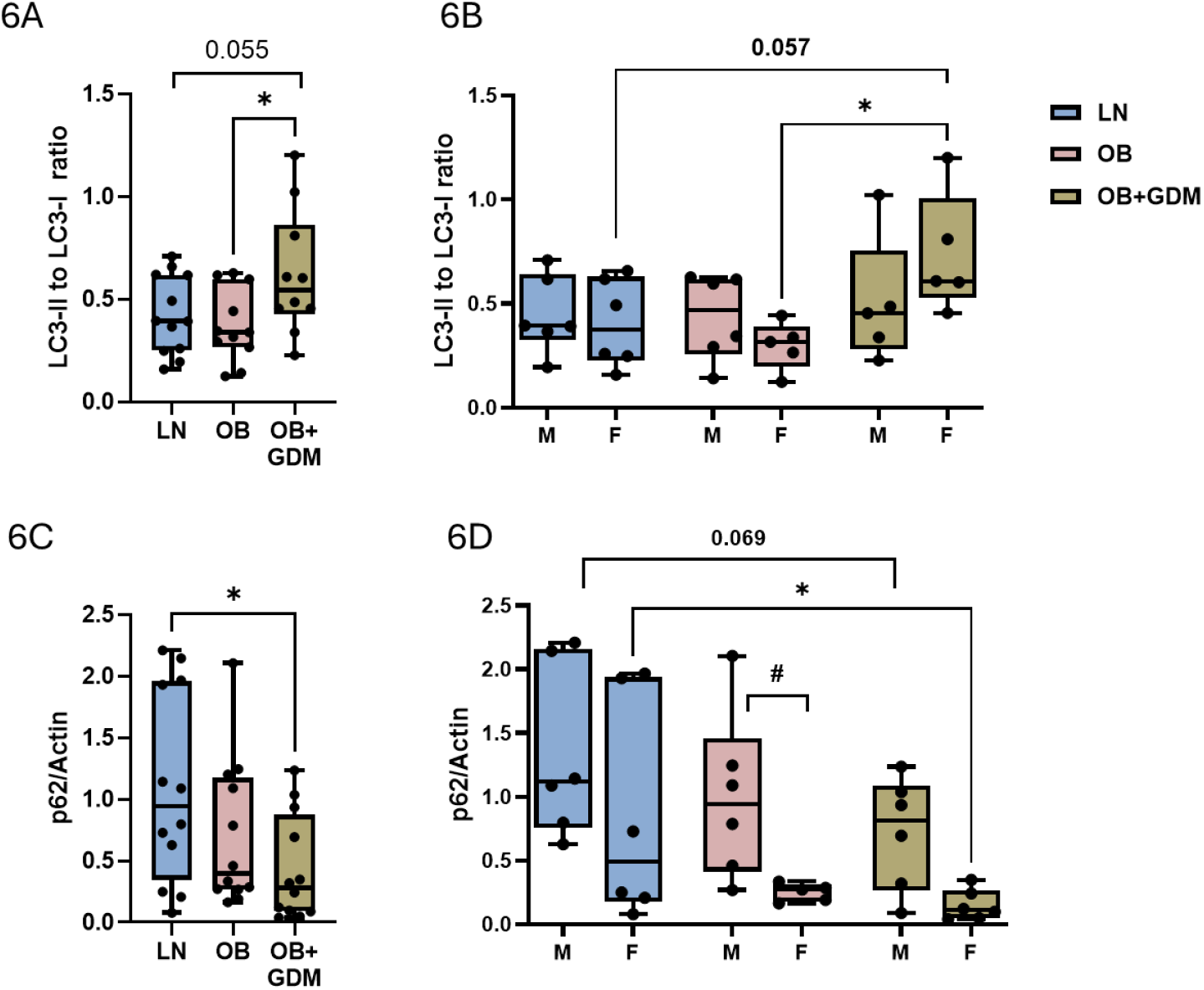
Expression of mitochondrial quality maintenance related proteins in villous tissue from LN, OB and OB+GDM placentas. A. B. Expression of LC3-II-I ratio and p62 in fetal sex combined and C, D in sex-stratified manner respectively. Data presented as box and whispers plot showing median, interquartile range, minimum and maximum values. * p<0.05.

## DISCUSSION

Mitochondria are indispensable for cellular and organ function including that of the placenta. A myriad of human conditions with different presentations such as Diabetes mellitus, Alzheimer’s disease, etc. can be traced back to dysfunctional mitochondria[28-32, 42, 43]. Mitochondrial content, morphology and their relative contribution to different cellular processes are determined by the specific demands placed on the cell. Several reports have provided evidence for mitochondrial dysfunction in placental pathologies including GDM[44]. However, most studies combine type A1 (dietary control) and A2 (requiring medication) GDM and compare against normoglycemic women which could lead to contradictory observations. One study distinguished between placentas from A1 and A2 GDM pregnancies using pregnancies in lean women as the control[44]. However, obesity is a major risk factor and increases the risk for developing GDM by 3 times. Here, we specifically focused on obese women with A2GDM as it presents the more severe phenotype[45-47]. We also included a group of women with obesity who did not develop A2GDM to further assess the individual effects of maternal obesity and obesity-associated GDM.

We observed an overall increase in signals for placental mitochondrial biogenesis, indicated by an increase in placental PGC-1α levels, with obesity but not OB+GDM when compared to the normoglycemic LN controls. A previous report showed reduced expression of PGC-1α in A2GDM placentas compared to LN controls, however that cohort also had significantly higher placental weights whereas in our study no differences in placental weight were observed between LN and OB+GDM groups, which could explain the contrasting results. This previous study also did not look for fetal sex-dependent changes. We observed that while male and female placentas from the LN and OB+GDM group had similar expression of PGC-1α, female placentas from the OB group had significantly higher levels compared to their male counterparts. PGC-1α is a master regulator of mitochondrial biogenesis that functions by increasing the expression of nuclear respiratory factors (NRF-1 and NRF-2), which in turn upregulate mitochondrial transcription factor A (TFAM). While we did not measure NRF1/2 levels, we found no changes in TFAM protein levels in obese placentas suggesting that the PGC-1α - TFAM axis might be dysregulated in obesity, suggesting distinct effects of maternal obesity on placental mitochondria. SIRT1 which acts upstream of PGC-1α was also increased (albeit not statistically significant) in obese placentas[48] suggesting that a slight increase in SIRT1 with obesity was sufficient to induce PGC-1α. PGC-1α is transcriptionally regulated by transcription factors such as PPARs, CREB, FOXO1, etc which may regulate its expression in placenta[49]. SIRT1 plays a crucial role in metabolism and is considered the master regulator linking cell metabolic status to transcriptional regulation of genes involved in lipid and glucose metabolism. Interestingly, studies in the non-pregnant state show downregulation of SIRT1 in peripheral tissues but an increase in hypothalamic SIRT1 associated with obesity highlighting its complex role in metabolism[50, 51]. Maternal obesity is associated with an increase in maternal plasma lipids and other metabolites, hence the increase in biogenesis stimuli could be interpreted as a placental response to maternal metabolic status. The distinct pattern of PGC-1α and SIRT 1 expression in obese placentas and absence of changes in GDM placentas (both male and female) imply that the maternal metabolic milieu in obesity and obesity-associated GDM affects placental mitochondrial response in a distinct complex manner.

Previously Abbade *et al* [44] reported a shift towards mitochondrial fusion in GDM placentae, which is in line with our observations. We see a significant decline in activated DRP1 and a trend towards an increase in OPA1 expression in GDM placentas. However, while activated DRP1 was significantly downregulated with obesity, OPA1 did not show any obesity-induced increase. Our fetal sex-stratified analysis showed that male placentas from the OB+GDM group had higher OPA1 levels compared to both LN and OB groups whereas the female placentas did not show this trend. OPA1 also showed sexually dimorphic expression in LN and OB placentas which was absent in GDM. We therefore conclude that the shift towards mitochondrial fusion in OB+GDM placenta is mediated by both increasing OPA1 and reducing activation of DRP1 whereas in obese placentas only DRP1 activation is regulated, potentially leading to disruption in mitochondrial dynamics in both groups. We also observed reduced expression of LONP1 in both OB and OB+GDM placenta. LONP1 plays an important role in mediating mitochondrial gene expression and protein quality control, by targeting misfolded proteins for degradation but also regulates mitochondrial dynamics by influencing both mitochondrial fusion and fission processes. TFAM binds to mitochondrial DNA and helps in mitochondrial DNA packaging as well as mitochondrial gene expression. Reduced expression of LONP1 in both obese and GDM placentas further suggests disruption of mitochondrial protein turnover and gene expression related activities[36, 52]. We believe this is reflected in the reduced expression of mitochondrial complexes observed in this study which could lead to lower mitochondrial oxidative phosphorylation and overall reduction in mitochondrial output. It is crucial to highlight that mitochondrial ultrastructure also plays an important role in determining spatial organization of these complexes in the inner mitochondrial membrane and by extension, their activity. While we did not assess mitochondrial ultrastructure in this study, based on previous reports on reduced mitochondrial function in trophoblast isolated from GDM placentae it could be implied that the reduced expression of complexes I-V in these placentae does reduce mitochondrial output[53].

Clearance of damaged cellular mitochondria via mitophagy is a crucial way of ensuring mitochondrial quality and function in cells. Under conditions of cellular stress, mitochondria undergo fragmentation by increasing fission and the fragmented mitochondria are then targeted for clearance. Our results suggested a shift towards fusion along with declining quality and function. We therefore assessed if this was exacerbated by the failure of placental cells to clear damaged mitochondria. Indeed, expression of PINK increased in obesity but there were no differences in proportion of activated PARKIN in any of the groups, suggesting that the mitophagy pathway was disrupted. As PINK labels damaged mitochondria for clearance, its increase in the obese group could be interpreted as attempts to remove damaged mitochondria which are thwarted by an inability to activate PARKIN. PINK1 senses mitochondrial damage by accumulating on the outer mitochondrial membrane (OMM) of depolarized mitochondria, triggering the recruitment of Parkin and initiating mitophagy. In healthy mitochondria, PINK1 is rapidly degraded after being imported into the inner membrane. However, upon mitochondrial depolarization, PINK1 accumulation is stabilized on the OMM resulting in recruitment of PARKIN, its phosphorylation and induction of the autophagy cascade. However, we did not observe any obesity-specific changes in the overall autophagy levels. The increase in LC3 flux ratio and reduced p62 with obesity and GDM suggested induction in autophagy as maternal condition worsened from obesity to GDM. We therefore conclude that in response to maternal obesity, the placenta attempts to increase mitochondrial biogenesis and mitophagy perhaps to rescue mitochondrial damage. This is reflected in the increase in SIRT1, PGC-1α, and PINK1. However, in response to GDM associated with obesity, these adaptations are not seen. Our results showing no changes in any of the above proteins in GDM coupled with reduced LONP1 expression suggests a selective decline in mitochondrial protein homeostasis and quality which could lead to the altered mitochondrial function reported in these placentas. Our results highlight (summarized in Fig 7) how maternal obesity and GDM have distinct effects on placental mitochondrial dynamics and clearance. Future studies on mitochondrial ultrastructure and function-based studies are needed to fully understand these distinct responses.

**Figure 7:**
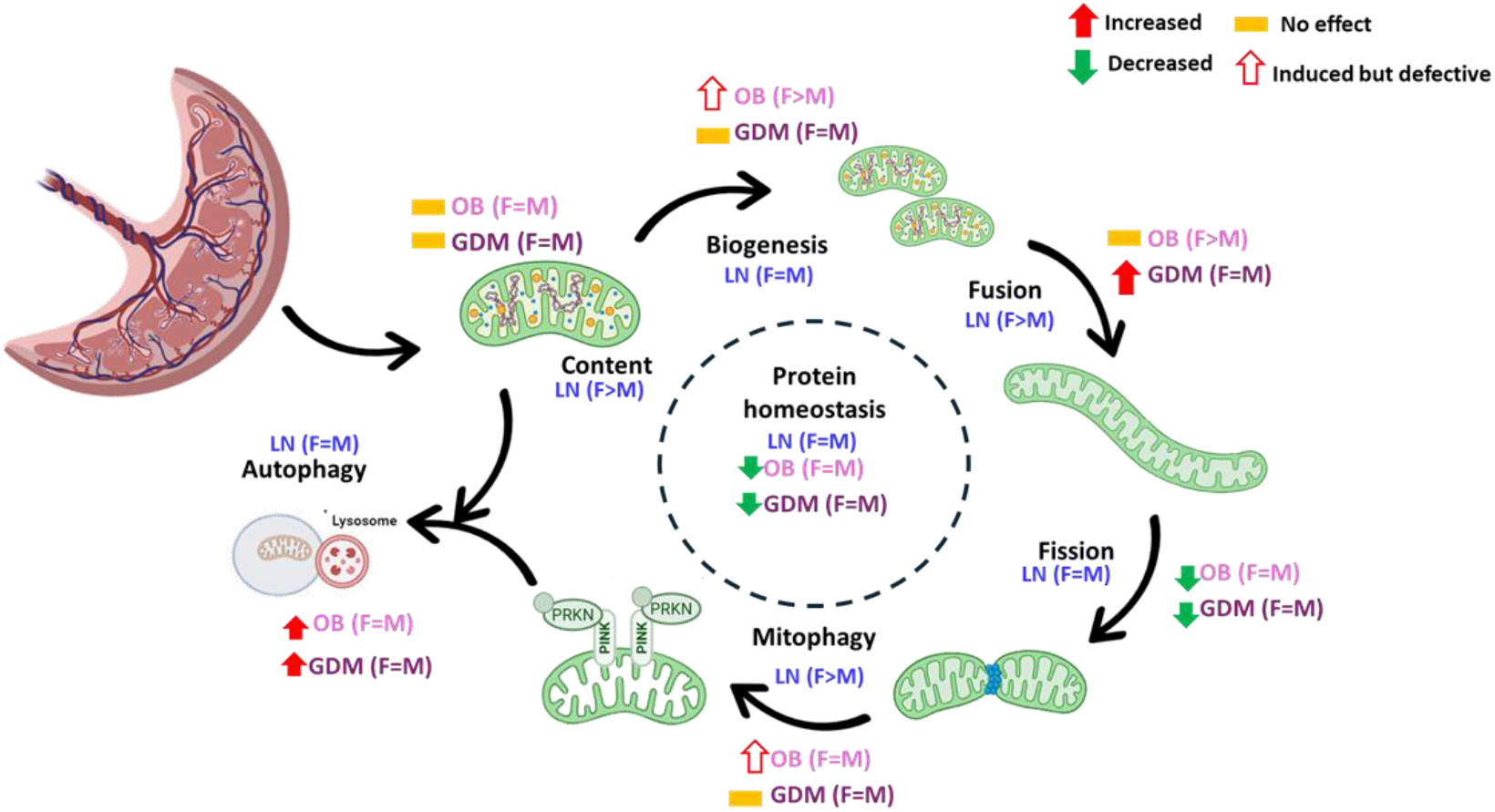
Overview of changes in placental mitochondrial dynamics due to maternal obesity and GDM.

## SUPPORTING INFORMATION

This article contains supporting information.

## AUTHOR CONTRIBUTIONS

LK conceptualized the study. KC and ET were involved in sample collection and with LK participated in data generation and analysis. LK and LM were involved with data interpretation and LK wrote the manuscript with contribution from LM.

