## Supplementary for "Dysregulation of placental mitochondrial structure dynamics and clearance in maternal obesity and gestational diabetes"

**Maternal obesity and Gestational diabetes alter placental mitochondrial dynamics in a sexually dimorphic manner.**

**Supplementary Table S1.0**

| Antibody | Host species, clonality | Company |
|---|---|---|
| Anti-SIRT1 | Rabbit, monoclonal | Cell signaling |
| Anti-PGC1a | Rabbit, polyclonal | Novus Biologicals |
| Anti-OPA1 | Mouse, monoclonal | Abcam |
| Anti-phosphorylated DRP1 | Rabbit, polyclonal | Cell signaling |
| Anti-DRP | Mouse, monoclonal | Santacruz |
| Anti-TFAM | Rabbit, monoclonal | Cell signaling |
| Anti-LONP1 | Mouse, monoclonal | Santacruz |
| Anti-p62 | Rabbit, polyclonal | Cell signaling |
| Anti-LC3 | Rabbit, monoclonal | Cell signaling |
| Anti – Beclin 1 | Rabbit, polyclonal | Cell signaling |
| Anti- OXPHOS | Cocktail of Mouse monoclonals | Abcam |

**SUPPLEMENTARY MATERIAL:**

**Supplementary figure S1**. Expression pattern of phosphorylated DRP (normalized to Actin) and Total DRP (normalized to Actin) in placenta from normoglycemic lean women (LN), women who are obese only (OB) and women with type A2GDM (GDM). **A,C**. sex combined manner, **B, D.** fetal sex stratified manner. Data expressed as Mean + SEM with n=6 per sex, per group.

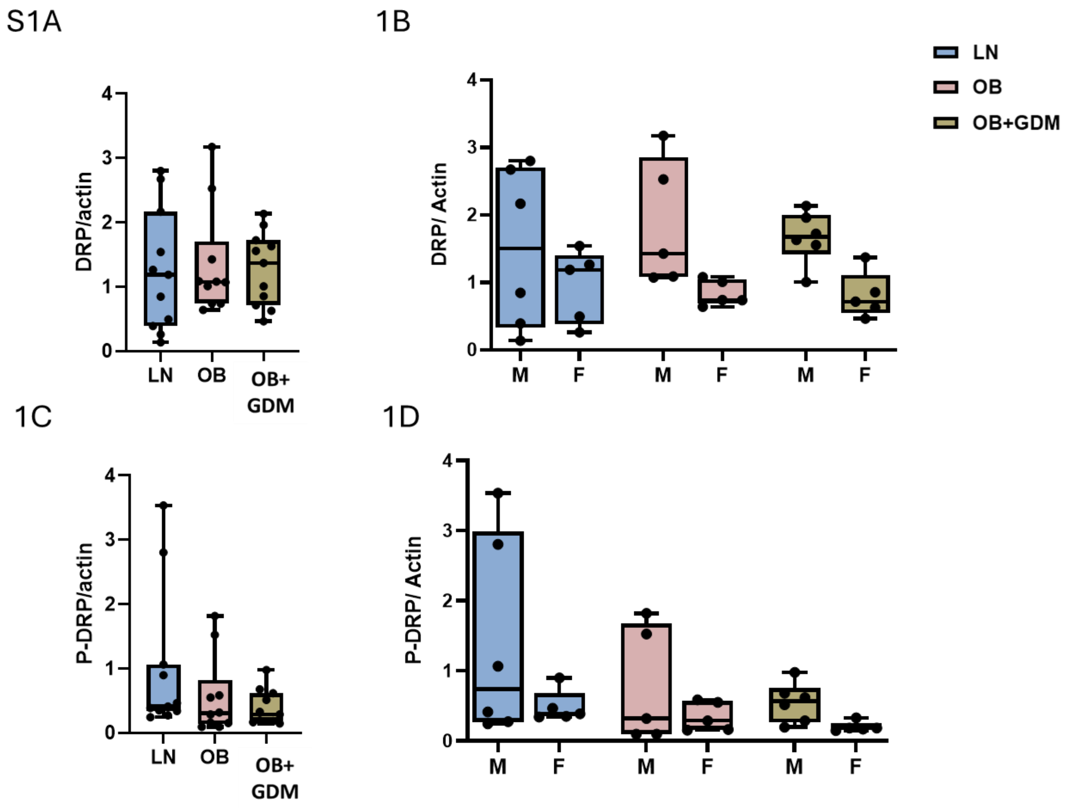

**Supplementary figure S2:** Expression pattern of phosphorylated PARKIN (normalized to Actin) and Total PARKIN (normalized to Actin) in placenta from normoglycemic lean women (LN), women who are obese only (OB) and women with type A2GDM (GDM). **A,C**. sex combined manner, **B, D.** fetal sex stratified manner. Data expressed as Mean <u>+</u> SEM with n=6 per sex, per group.

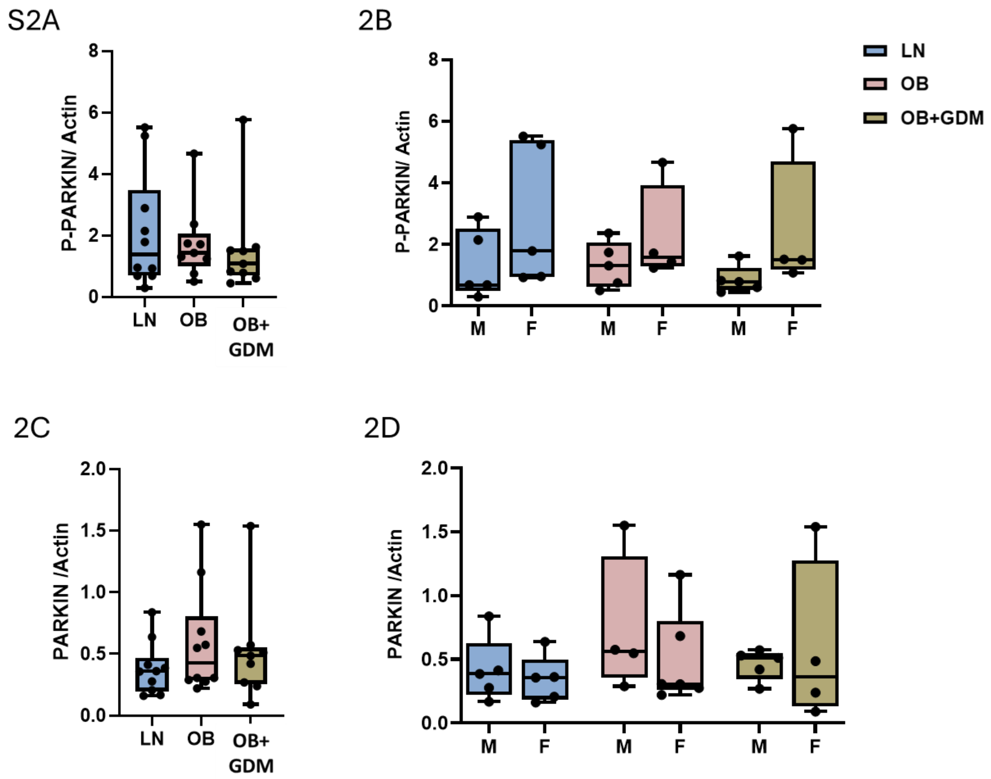

**Supplementary figure S3:** Expression pattern of LC3-I (normalized to Actin) and LC3-II (normalized to Actin) in placenta from normoglycemic lean women (LN), women who are obese only (OB) and women with type A2GDM (GDM). **A,C**. sex combined manner, **B, D.** fetal sex stratified manner. Data expressed as Mean + SEM with n=6 per sex, per group.

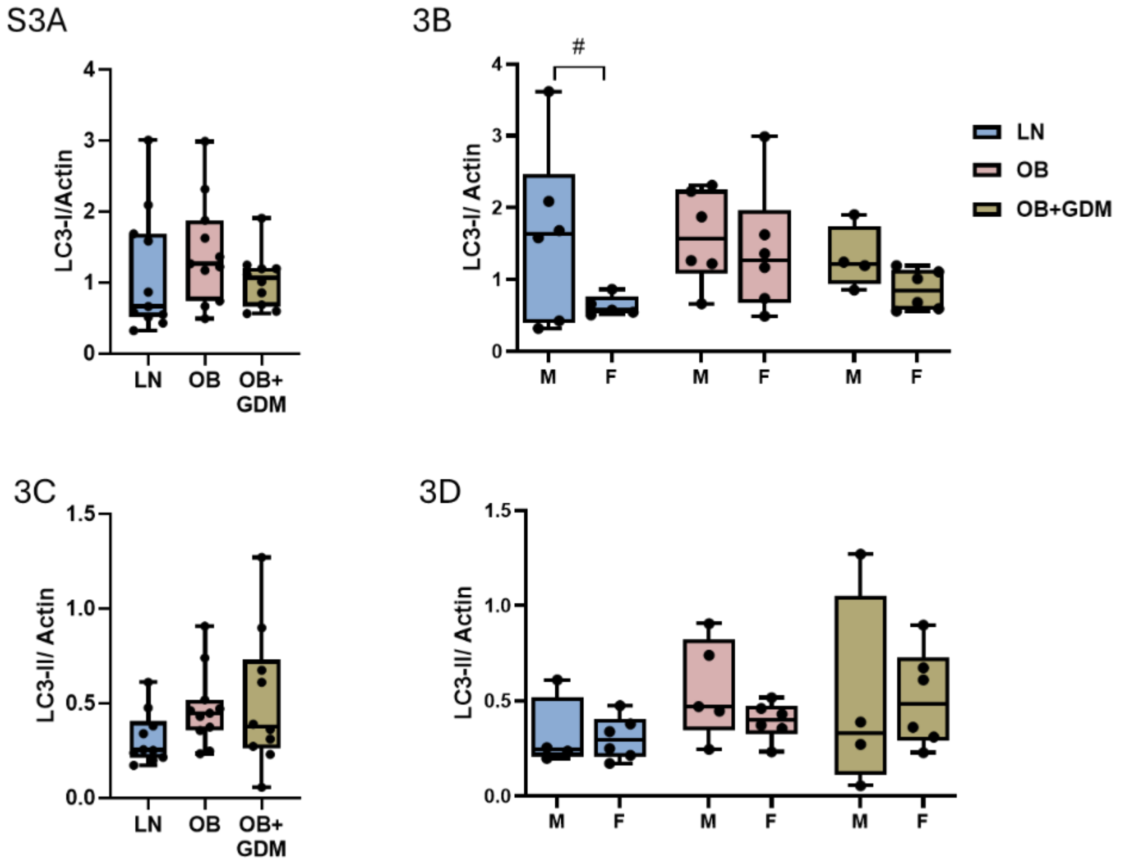
